# Contributions of source populations, habitat suitability and trait overlap to benthic invertebrate community assembly in restored urban streams

**DOI:** 10.1101/2024.07.01.601525

**Authors:** Svenja M. Gillmann, Nele Schuwirth, Armin W. Lorenz, Daniel Hering

## Abstract

Community development in restored streams is often slow or even absent, but reasons remain obscure. Inadequate restoration measures, catchment-scale pressures, community closure and colonization barriers all may prevent or slow down recovery processes. For initial colonization, dispersal processes are supposed to be most important, which are referred to as dispersal filter. Environmental conditions of a restored reach determine if a dispersing species can successfully establish (environmental filter). Lastly, while available niches at those reaches fill up, biotic interactions, such as competition, become more important (biotic filter). To investigate the importance of these different filters, we compared benthic invertebrate communities of 20 sites in the Boye catchment (Western Germany), a former open sewer system. The sites were grouped, based on the years since restoration, into ‘unimpacted’ (never restored), ‘recently restored’ (< 4 years) and ‘mature restored’ (> 10 years) sites. Data collected at 28 additional sites in the catchment informed us on distances to potential source populations. Habitat suitability describes the fit between environmental conditions (abiotic site data) and species preferences and was used to assess the role of environmental filtering. We evaluated the role of the biotic filter based on trait overlap, referring to possible interspecific competition.

Communities collected at recently restored sites differed from those of mature restored and unimpacted sites. Taxa present at recently restored and mature sites had closer source populations than those of unimpacted sites. Taxa at mature and unimpacted sites had a better fit to the present habitats than those of recently restored sites. The trait overlap did not differ between co-occurring and not co-occurring taxa at any of the site groups. Our findings show that communities of mature restored sites that have been restored more than 10 years ago, resembled those of unimpacted sites. Dispersal was most important in early years of recovery. Taxa occurrences at sites with nearby source populations and low habitat suitability are likely the result of high rates of dispersal from upstream sources (mass effects). These can be caused by hatching events or environmental disturbances. Habitat suitability played a larger role for communities at mature and unimpacted sites which indicates that optimal communities shape over time. We did not find indications that competition played a role for community assembly. Hence, dispersal and habitat suitability were most relevant for species’ occurrences. Competition could be more important on micro scales and the results may differ if species abundances are taken into account.

**Graphical Abstract:** 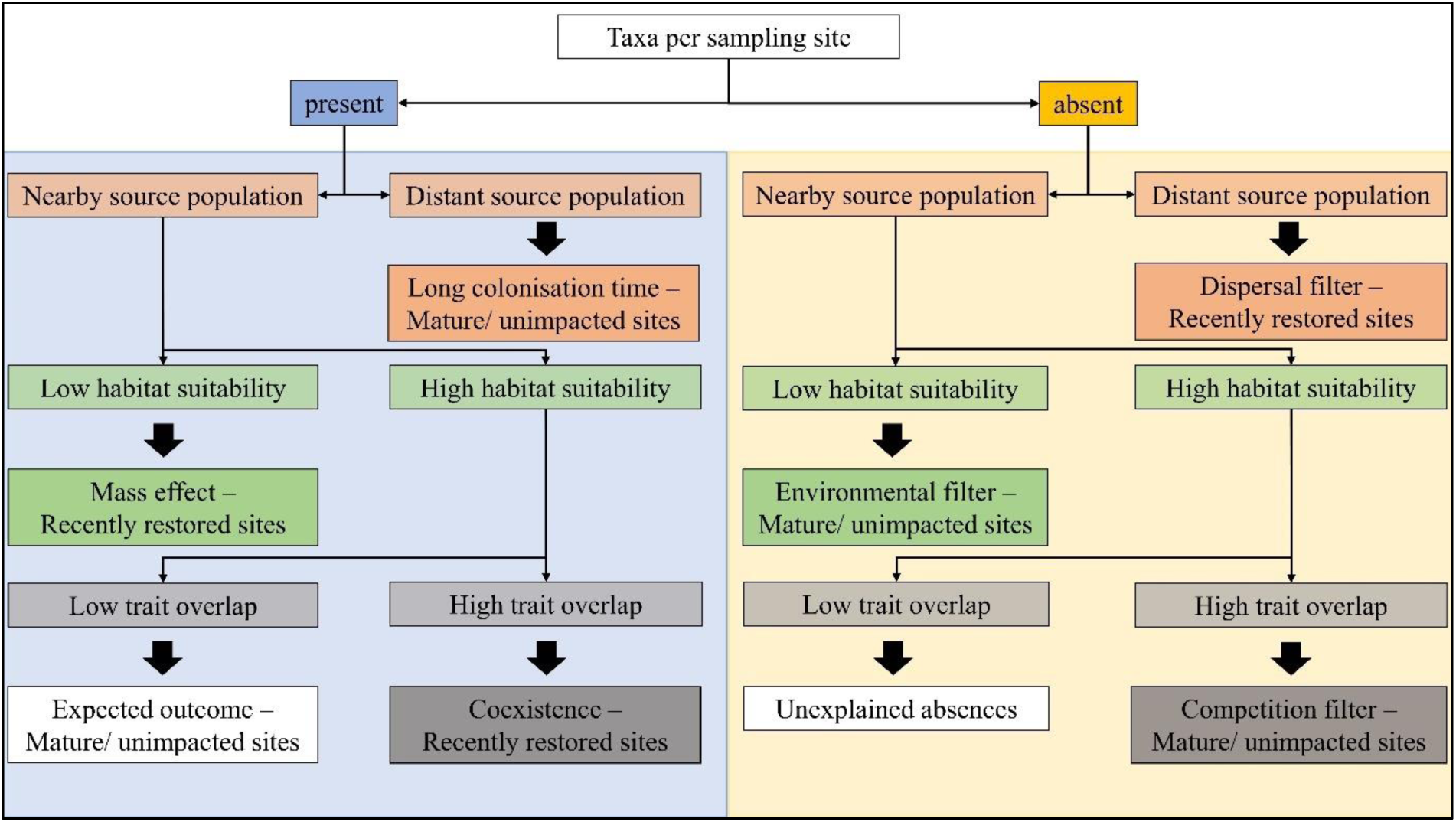

## Introduction

The recovery of aquatic communities after pressure release can take multiple pathways (Sarr, 2002; Vos et al., 2023). These range from persistence of the degraded community (Lorenz et al., 2018) to full recovery (Clements et al., 2021). The degree of recovery depends on successful colonisation of a diverse community. However, even if communities recover at first, new and/or persisting pressures can interrupt and stop the progress of recolonisation (Haase et al., 2023). Understanding the processes that govern community assembly, is crucial to improve planning of future restoration measures and efficiently support stream communities on their way to full recovery.

From a more theoretical viewpoint, recovery processes are closely related to metacommunity theory. Metacommunity assembly is governed by spatial and local processes (Leibold et al., 2004; Lake et al., 2007; Vos et al., 2023). The dispersal ability of a species, the distance to source populations and dispersal barriers represent the first filter that determines whether a restored reach will be colonised (Lake et al., 2007; Sundermann et al., 2011; Tonkin et al., 2014). Consequently, good dispersers are the first to reach those sections (Winking et al., 2016; Baumgartner & Robinson, 2017; Lorenz et al., 2018), which is expected to be most relevant in the starting phase of community recovery (Vos et al., 2023). However, dispersal capacity loses importance with increasing connectivity (Brederveld et al., 2011). Recolonisation is particularly strong when source populations are in close proximity to the recovering stream reach and mass effects occur ( Leibold et al., 2004; Heino et al., 2015; Brown et al., 2018; Tonkin et al., 2018). These describe the phenomenon that species occur in unsuitable habitats, due to high rates of recurring dispersal from nearby source populations (Leibold et al., 2004).

The environmental filter, or suitability of a given habitat, eventually determines whether the colonising species can establish a population (Lake et al., 2007). Streams provide a variety of habitats and thus ecological niches which are colonised by aquatic biota according to their preferences (Schmera et al., 2017). The ecological niche is defined by the range of conditions relating to water quality, preferred habitat and food source (Soberon & Peterson, 2005). The niche theory distinguishes between the fundamental niche, which defines the conditions that are generally suitable for a given species and the realised niche, where the species actually lives under the conditions of competition and other biotic interactions (Soberon & Peterson, 2005). Aquatic communities therefore react sensitively to changes in the environmental conditions and stressed systems are only inhabited by tolerant species (Rumschlag et al., 2023). Thus, species tolerances against stressors are expected to be less important if restoration measures have reduced stress intensities (Vos et al., 2023). Within established communities, local dynamics are governed by biotic interactions, e.g. competition, which importance increases over time (Lake et al., 2007; Vos et al., 2023). Regional community dynamics are shaped by an equilibrium between local extinctions and colonisations, which can be driven by random processes (neutral theory) ( Leibold et al., 2004; Larsen et al., 2018) or short-term disturbances.

These processes acting during community assembly are closely intervened, thus, it is important to analyse their role for recolonisation simultaneously. However, while the processes have been analysed individually or in pairs, studies encompassing all three filters are lacking (Liu et al., 2023). Most studies focus on the influence of environmental and dispersal filters, while the role of biotic interactions is often neglected (Heino et al., 2015; Tolonen et al., 2018; He et al., 2023; Zheng & Yin, 2023).

Among riverine organisms, benthic invertebrates are an ideal group to test metacommunity theory. They are species-rich, their ecological preferences are well understood, and they contribute to a range of functional processes of streams, e.g., decomposition of organic matter (Palmer & Poff, 1997; Schmera et al., 2017). Benthic invertebrate species differ in sensitivity toward environmental changes, dispersal capacity and ecological niches (Jowett & Richardson, 1990; Kenney et al., 2009). Their specific preferences for environmental conditions can be used in habitat suitability models, to predict their potential occurrence in stream networks (Hirzel & Le Lay, 2008) and analyse changes in community composition (Lee et al., 2023), Recently, species preferences were fed into the models as prior knowledge, to improve the performance of the models (Vermeiren et al., 2020, 2021). However, integrating dispersal and biotic interactions into habitat suitability models is difficult, and has rarely been done (Schuwirth et al., 2016).

For a detailed assessment of the three community assembly filters (dispersal, environment, biotic interactions), extensive monitoring of the recovering communities is needed. Previous studies revealed that a benthic invertebrate community needs at least eight years to reach a certain maturity after restoration ( Lorenz, 2020; Gillmann et al., 2023) . Depending on the size of the restoration measures, new habitats have to develop first, before a stable community can establish (Pilotto et al., 2022). Further, restored sites were shown to be more rapidly colonised, if source populations are within close proximity (Sundermann et al., 2011; Tonkin et al., 2014). Vos et al. (2023) predicts the role of dispersal, tolerance and competition to change over time, with dispersal being most important in the first phase of recolonisation, before environmental filters and biotic interactions gain relevance. These predictions were generally supported by Gillmann et al. (2024): The analysis of 10-year post-restoration benthic invertebrate community assembly revealed that dispersal capacity and tolerance toward organic decomposition decreased with time since restoration, while interspecific competition increased. While this previous study revealed how the three filters generally change over time at the community level, it does not provide a species- and site-specific analysis. The environmental filter was only analysed in terms of tolerance toward chloride and organic decomposition, without considering substrate availability and other abiotic factors. Additionally, mean community dispersal capacity only gives limited information on the presence of a dispersal filter.

Here, we used data of benthic invertebrate communities, collected at specified distances, to gain information on potential source populations for our ‘main sites’. To investigate the role of the environmental filter, we matched the abiotic site data (‘site profiles’) with the species’ preferences (‘species profiles’), which resulted in a measure of habitat suitability (Vermeiren et al., 2020). Trait overlap served as a proxy for biotic interactions. With this detailed dataset we compared how the role of the three filters differs between sites of different maturity stages. More specifically, we hypothesized that (1) the community composition of recently restored sites differs from those of mature and unimpacted sites, as new habitats are still developing and not all potential taxa reached the restored sites. (2) The closer the nearest source population, the more likely is a species’ occurrence. This is particularly relevant for recently restored sites, and less for mature and unimpacted sites. (3) The better the ‘species profile’ matches the ‘site profile’, the more likely is its occurrence. This ‘environmental filtering’ is supposed to be less relevant for the recently restored sites, but most relevant for mature and unimpacted sites. (4) The more strongly traits of a given species overlap with species established at a site, the less likely is its occurrence. This biotic filtering is particularly relevant for mature and unimpacted sites.

This study aims at a deeper understanding of metacommunity assembly, by capturing the major assumptions of the Asymmetric Response Concept (ARC; Vos et al., 2023) in detail. The comparison of different maturation stages of restored sites will help to inform managers, which actions are most important during the respective phases of recovery.

## Methods

### Sampling

The Boye is a tributary to the Emscher catchment, located in Western Germany. The Boye and most of its tributaries were used as open sewers for several decades. At the end of the 20^th^ century, the sewage system was moved underground and the impacted streams were gradually released from wastewater, followed by hydromorphological restoration. Restoration measures at the last stream reaches were finalised in 2021. A more detailed description of these measures and the Boye catchment itself is provided in Gillmann et al. (2023).

In total, 48 sites were sampled for their benthic invertebrate community. Twenty of the sites are part of an annual monitoring program, established for the RESIST project (‘main’ sites in Fig. 1). These were grouped according to the time of their completed restoration into ‘unimpacted’ (never carried wastewater), ‘mature restored’ (restored > 10 years prior sampling) and ‘recently restored’ (restored < 4 years prior sampling). To gain information on potential source populations for these sites, additional 28 sites (‘source’ sites in Fig. 1) were distributed in the catchment and sampled in 2022. These sites were placed 1 km and 2 km upstream of each main restored site. If possible, an additional site was located in the upstream sections of the streams that never carried wastewater. The unimpacted ‘main’ sites were considered as potential ‘source’ sites; hence, no additional sites were sampled upstream of these if they were within the 2 km distance of another main site.

**Figure 1:**
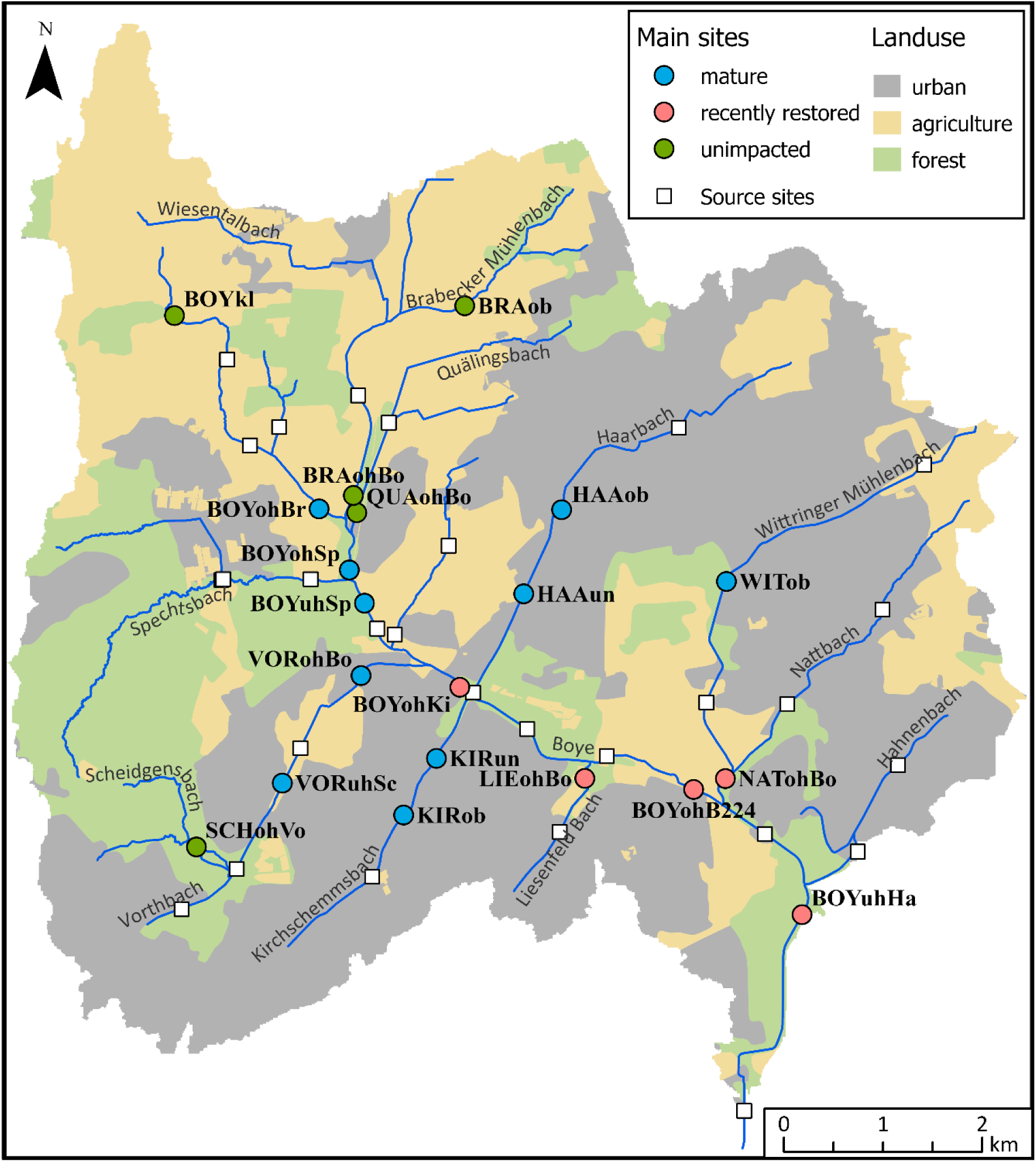
Map of the Boye catchment (Ruhr Metropolitan Area, Western Germany). The ‘main’ sites are displayed with dots, colored according to their site group: mature (blue), recently restored (red), unimpacted (green). The ‘source’ sites, located 1 and 2 km upstream of each main site, are displayed as white boxes. Three major types of land use of the catchment (merged from CORINE and InVeKos) are displayed in different colors: urban (grey), agriculture (yellow), forest (green).

All benthic invertebrate samples were collected following the standardised protocol for multi-habitat sampling (Haase et al., 2004). The collected samples were preserved in ethanol (96%) and brought back to the laboratory for further processing. The ‘main’ sites are part of an annual sampling campaign. To comply with the methods used in previous years, these samples were sorted and identified morphologically, according to the operational taxalist for Germany (Haase et al., 2006), resulting in a list with species abundances.

Standard methods for monitoring of benthic invertebrates (Haase et al., 2006) provide reliable information about abundances but are time consuming. Alternative methods based on eDNA and DNA metabarcoding techniques (Elbrecht et al., 2017) allow high throughput but provide only presence/absence information. Hence, we used DNA metabarcoding to identify the samples of the ‘source sites’ (Buchner et al., 2021) and adjusted them to the same taxonomic level as the ‘main’ sites.

### Environmental data

At all sites, the proportions of available stream bottom substrate were recorded in 5% increments, as specified in the standardized protocol for MHS (Haase et al., 2004). Single measurements of oxygen, temperature, conductivity, and pH were recorded for the ‘main’- and ‘source’ sites. More detailed physico-chemical parameters were collected only at the ‘main’ sites: Upon MHS, flow velocity was measured at each location of sample unit collection and summarised into a mean flow velocity per site. Additionally, two loggers per site recorded water temperatures at 30-minute intervals. We used the data collected from June until September 2021, i.e., the year before sampling of benthic invertebrates, to determine mean summer water temperatures. Lastly, water samples were collected bi-weekly, starting in March 2021, which were stored at -20 °C before analysis commenced in the laboratory. Concentrations of ammonium, nitrate, dissolved oxygen content (DOC) and orthophosphate were derived from the water samples. In combination with the oxygen content, measured on site, they were used to determine the water quality classes related to saprobity.

### Data preparation

#### Distance to source populations

We calculated the terrestrial and instream distance between the 48 sampling sites. Only distances to the upstream sites were considered for further analysis. Taxa were grouped into holo- and merolimnic taxa. For every combination of taxon and site, we determined the terrestrial and instream distance to the closest source population, meaning the next upstream site with an occurrence of the taxon. For hololimnic taxa, only the instream distance was considered, while for merolimnic taxa, the smaller of the instream and terrestrial distances was considered as closest distance to the next source population.

#### Environmental filtering

We analysed the role of environmental filtering for the 20 ‘main’ sites by calculating habitat suitability values per species and site. To this end, we compared ‘species preference profiles’ with the environmental conditions at each of the ‘main’ sites: substrate proportions, mean flow velocity, mean summer temperature and saprobic classes. For the species preference profiles, the following parameters were collected: microhabitat preferences, saprobic values and indicator weights, KLIWA index and specificity value (freshwaterecology.info; Schmidt-Kloiber & Hering, 2015) and flow velocity preference (STOWA database; Verberk et al., 2012).

Habitat suitability was derived for the parameters, ‘substrate’, ‘saproby’ and ‘temperature’ for each combination of site and species. To compare the present substrates with the taxa preferences, some of the substrate categories were combined to fit the categories of substrate preferences as defined in freshwaterecology.info (Appendix S1, Table S1). The taxas’ saprobic values and indicator weights were compared to the saprobic classes of each of the sampling sites as derived from water quality parameters. Saprobic classes were calculated based on the class borders defined by the LAWA, in 1998, published by Bernatowicz et al. (2009)see also Appendix S1, Table S2). For the calculations, the data collected from the bi-weekly water samples in 2021 was used. The following parameters were included in the calculation: minimum oxygen contents, maximum ammonium, maximum nitrate, total ammonium and median nitrate, mean DOC, maximum and median orthophosphate. For water temperature, we used the logged temperatures from summer 2021 (June until September), as the corresponding biotic metric (the KLIWA index) is based on the summer mean and maximum temperatures.

The habitat suitability functionsquantify how the habitat’s suitability for each taxon varies with each environmental variable, scaling from zero to one. They consist of preference scores, which are normalized to values between zero and one, and a linear interpolation to derive a continuous function from discrete classes of the environmental variable (see Appendix S1, Table S3). This approach has previously been used by Vermeiren et al. (2020), where the habitat suitability served as prior knowledge input in habitat suitability models. Instead of using a full habitat suitability model, which would require a larger number of sites for calibration, we used the habitat suitability functions directly to predict species occurrences and absences. To this end, we calculated the mean habitat suitability from all environmental factors.

#### Biotic interactions

The resource use of benthic invertebrates is mainly defined by its feeding type and microhabitat preference. Therefore, we utilized these two key traits to analyse biotic interactions between taxa in terms of trait overlap. After downloading the trait information from freshwaterecology.info (Schmidt-Kloiber & Hering, 2015), we quantified the trait overlap using the ‘gawdis’ function (Bello et al., 2021), resulting in a Gower Similarity index, spanning between 0 and 1, for each species pair. We defined the highest possible match between traits of different taxa as the degree of possible competition between them. Therefore, we determined the maximum trait overlap per taxon for co-occurring or not co-occurring taxa.

#### Data analysis

We calculated the Jaccard dissimilarity between communities, based on the presence/ absence taxalist of the ‘main sites’, using the function ‘vegdist’ (package ‘vegan’, v.2.6-4, Oksanen et al., 2024). With the function ‘metaMDS’ (package ‘vegan’, v.2.6-4, Oksanen et al., 2024), we visualised the community distances using Non-metric multidimensional scaling (NMDS, k=3). The site labels include the stream, where the site is situated (first three letters), and the description of the location within the stream (fourth letter onward). The difference between the different site groups (‘unimpacted’, ‘recently restored’, ‘mature restored’) was tested, using the Analysis of Similarities (ANOSIM) statistics (function ‘anosim’, package ‘vegan’, v.2.6-4, Oksanen et al., 2024).

We separately analysed each of the three factors (dispersal, environmental filtering and biotic interactions) and compared values between the present and absent communities. To visualize changes in the proportions of present and absent taxa along the gradients of the different filters, we summarised the number of cases for subgroups along these gradients. The distance to the nearest source was used to quantify the dispersal filter. Due to our sampling design, most sites were 1 km apart, however, some ‘main’ sites were closer together. To consider these smaller scales, we divided the groups in 500 m steps, resulting in 18 groups. Cases, where taxa had no upstream source population were grouped together as ‘no source’ (19^th^ group). The mean habitat suitability and maximum trait overlap, both ranging from zero to one, were divided into 10 groups. For each group per factor, we calculated the proportions of present vs. absent taxa. In addition, we determined the total number of cases, contributing to each proportion per subgroup.

We used logistic regression models to analyse the probability of species occurrence per factor for all sites. Therefore, we constructed binomial generalised linear mixed models (GLMMs) with ‘logit’ link function (package ‘glmmTMB’, v.1.1.8, Brooks et al., 2017), with occurrence as binary response variable. We used the factors (distance to nearest source population, mean habitat suitability, trait overlap) as fixed effect, respectively, and included the species as random effects. The conditional R-squared results from correlating the fitted with the predicted values. We checked the residuals, using the ‘simulateResiduals’ from the ‘DHARMA’ package (v.0.4.6, Hartig & Lohse, 2022).

For each factor, we analysed the difference between groups of occurrence and groups of sampling sites (i.e., ‘unimpacted’, ‘recently restored’ and ‘mature restored’ sites). The spread of the data was visualised using boxplots. To statistically determine the difference between groups, we constructed different models per factor. For the distance to the nearest source, we used a GLMM with gamma distribution (package ‘glmmTMB’, v.1.1.8, Brooks et al., 2017). For habitat suitability and trait overlap, we used generalized additive models with beta and beta one inflated distributions (package ‘gamlss’, v. 5.4, Rigby & Stasinopoulos, 2005). All models treated ‘occurrence’ and ‘site group’ as fixed effects and ‘species’ as random effect. We extracted the estimated marginal means (EMMs) from the models using the function ‘emmeans’. From the same package, we used the function ‘pairs’ to determine the significance of differences between present and absent taxa per group, which conducts a post-hoc test between the EMMs of different groups, while considering unequal number of observations and random effects (package ‘emmeans’, v. 1.10.2, Lenth, 2017).

All data was analysed in R (v.4.1.2, R core Team, 2021) and visualised with the package ‘ggplot2’ (v.3.4.1, Wickham, 2016).

## Results

### Description of the benthic invertebrate communities

In total, we identified 107 different taxa at the main sites in the Boye catchment. Individuals belonging to the family of Chironomidae occurred at all main sites. The taxa identified to species level with the highest prevalences were *Gammarus pulex*, *Limnephilus lunatus* and, *Prodiamesa olivacea*. They occurred at 18 of the 20 main sites. Our dataset included 34 taxa that each occurred at only one of the main sites. 37 taxa were only found at source sites.

### Differences between benthic invertebrate communities

The NMDS (Fig. 2, stress= 0.13) displays the Jaccard similarity between communities of the main sites. The communities of the mature and unimpacted sites are clustered close together, displaying their high degree of similarity. Three of the recently restored sites are clearly apart from the other communities, while one of the recently restored sites is more similar to the unimpacted and mature sites than to other sites of its group. The Analysis of Similarities (ANOSIM) confirmed a moderate difference between groups of sampling sites (R²= 0.22, p= 0.02).

**Figure 2.**
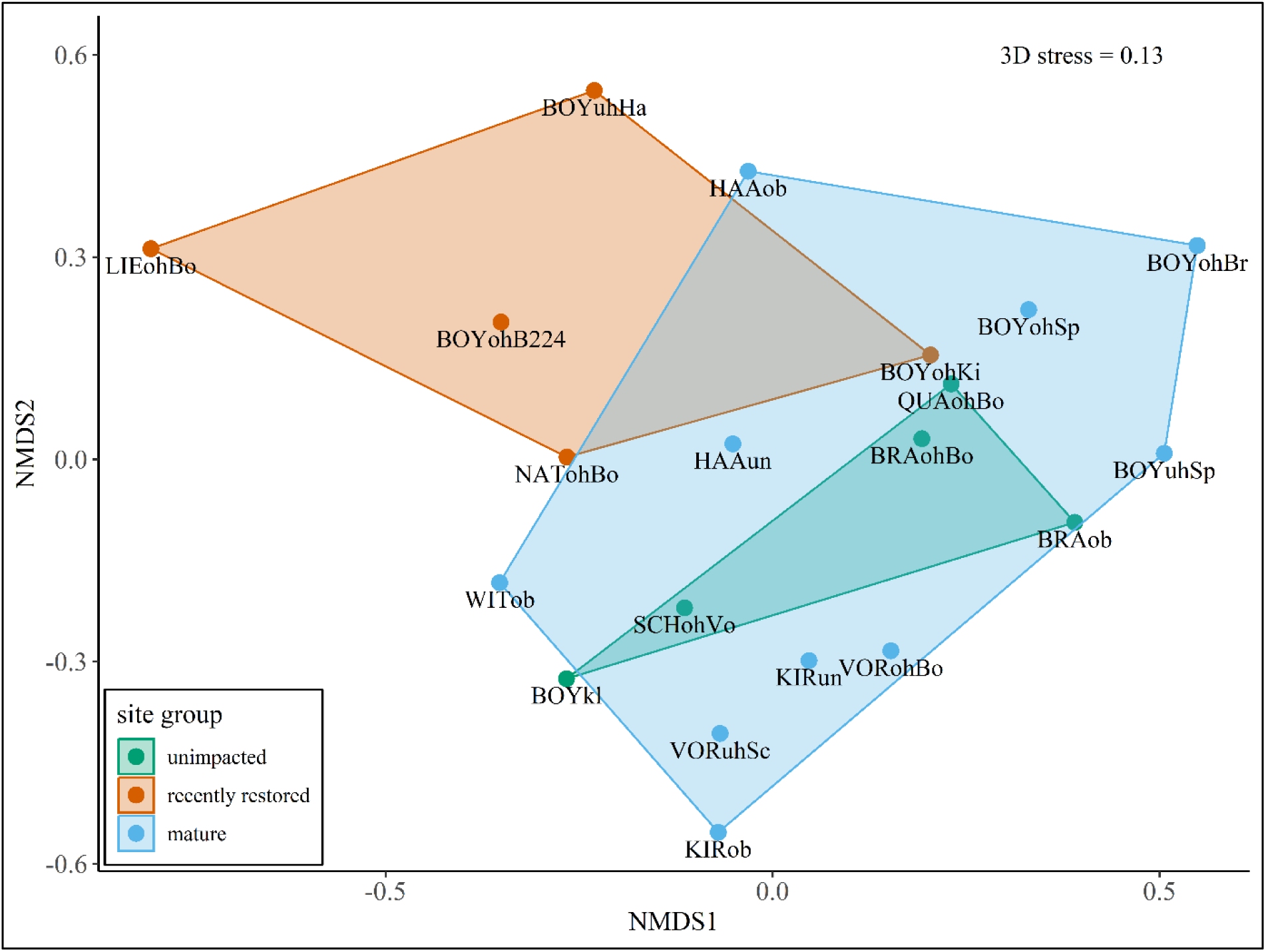
NMDS of Jaccard dissimilarities, derived from presence/ absence data of taxa found in the different sites (stress = 0.13). Differences between site groups were tested using the ANOSIM statistics (R²= 0.22, p= 0.02).

### Dispersal filter

We determined the distance to the nearest source population for all taxa per site and analysed the changing proportions of present and absent taxa along the distance gradient. The proportion of present taxa decreases with the distance to the nearest source population (Fig. 3A). This pattern is supported by the underlying model which identifies a significant decrease in probability of taxa occurrence (p < 0.05, Fig. 3B) with increasing distance to the source. In the majority of cases, a source population is present within 1.5 km upstream. Only in one case, a taxon was found to occur with its nearest source population located at a distance > 5 km. In a large proportion of cases, absent taxa did not have an upstream source population within the Boye catchment which was true for fewer cases of present taxa.

**Figure 3.**
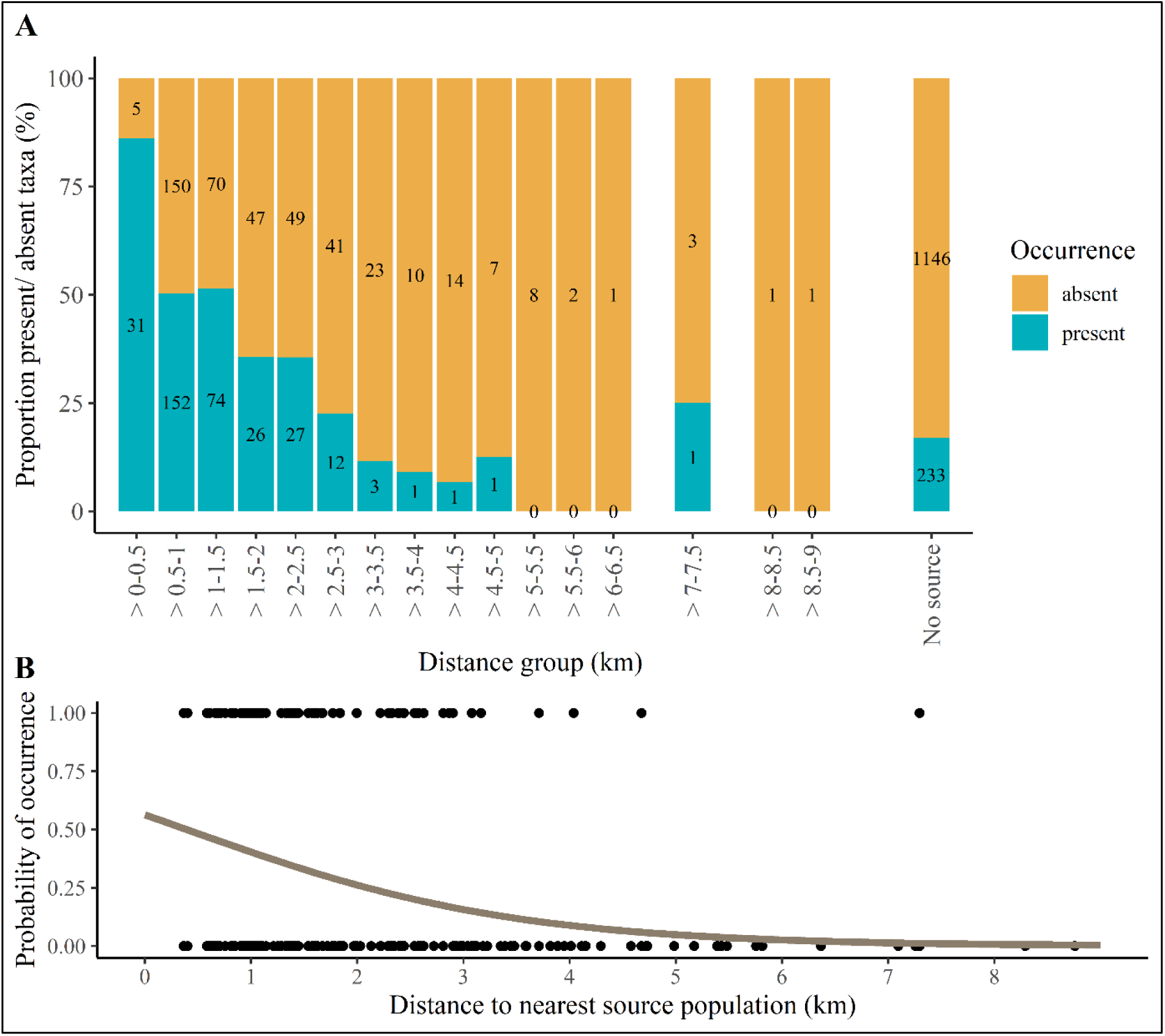
A: Proportion of present and absent taxa per distance group. The distances were discretized into classes of 500 m, resulting in 18 groups. Taxa that did not have an upstream source population were defined as 19^th^ group. The number of ‘absent’ and ‘present’ cases is indicated in the corresponding bars. B: Logistic GLMM displaying the probability of occurrence in relation to distance to the nearest source population (R² = 0.27).

We determined the distribution of the distances to the nearest source population across unimpacted, recently restored and mature sites, categorized by ‘absent’ and ‘present’ taxa for each site group (Fig. 4). Statistical differences between groups were determined using estimated marginal means. At unimpacted sites, there is no significant difference in distance between ‘absent’ and ‘present’ taxa (p = 0.152). The distance to the nearest source population is significantly different between ‘absent’ and ‘present’ taxa for both, the recently restored and the mature restored sites (p < 0.05). In both cases, the distance to population sources is lower for the present taxa. However, the distribution of the data and estimated marginal means are larger for absent taxa at the recently restored sites compared to the mature sites.

**Figure 4.**
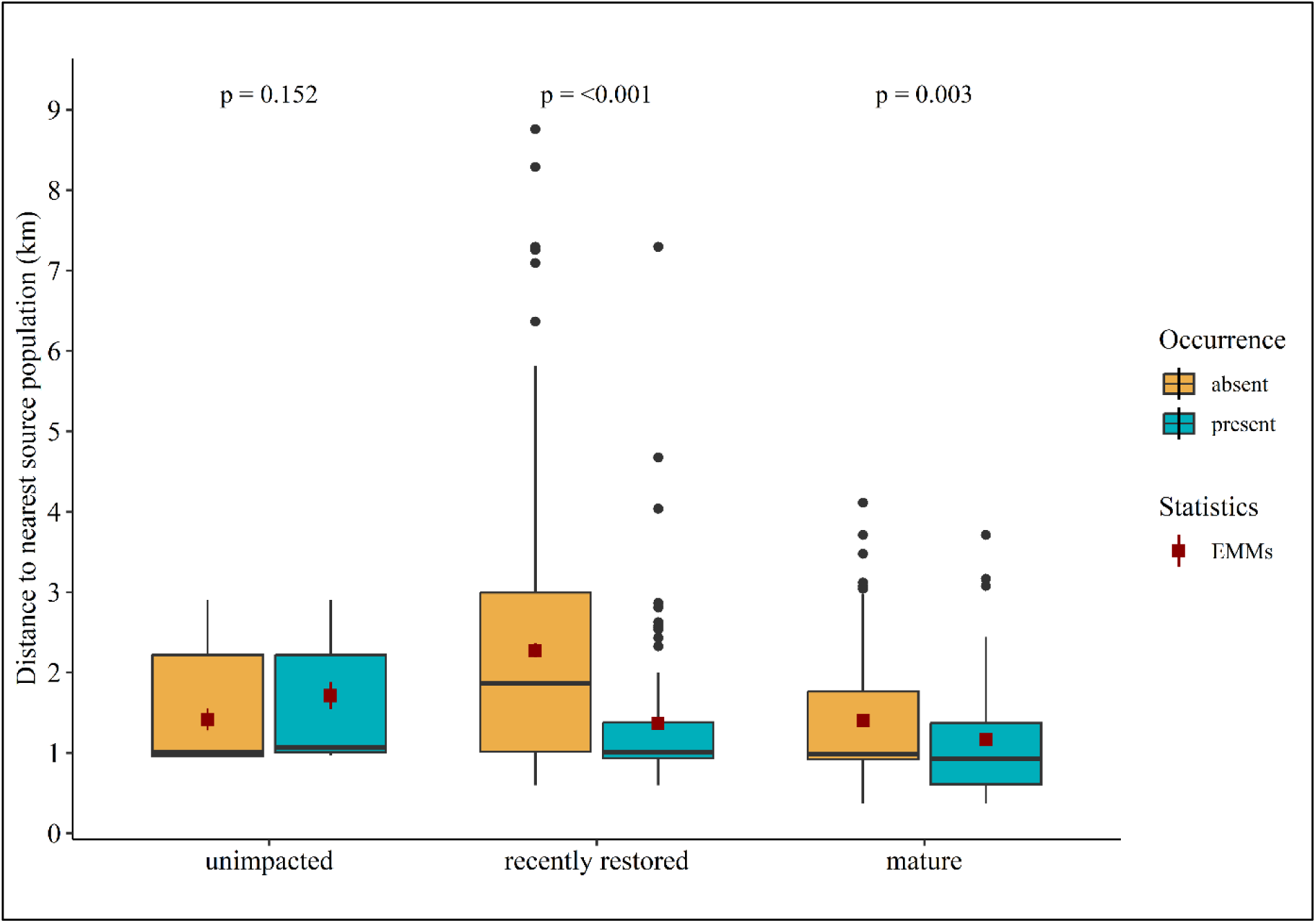
Boxplot of present and absent taxa and their distance to the nearest source population, grouped by maturation state. The difference between distances of the absent and present taxa were compared conducting a post-hoc test on estimated marginal means (EMMs). These were extracted from the corresponding generalised linear mixed model (GLMM), to account for different numbers of observations and to consider random effects. Estimated marginal means (EMMs) are represented by red boxes and corresponding error bars. Non-overlapping error bars indicate statistically significant differences (p < 0.05). For more detailed results on EMMs see Appendix S2, Figure S1.

### Environmental filter

We used the mean habitat suitability to compare differences between present and absent taxa per site. The proportion of present species in all groups is below 50%, except for the group with the highest habitat suitability (Fig. 5A). The group with the lowest habitat suitability has the lowest proportion of present taxa. In the habitat suitability classes between 0.1 and 0.5 the proportion of present taxa varies around 25% and increases with increasing habitat suitability above 0.5. This increasing trend is mirrored in the modelled probability of taxa occurrence (p < 0.05, Fig. 5B). However, the two classes with the highest habitat suitability have the lowest number of cases (73 and 14). The largest number of cases belong to the group with mean habitat suitabilities between 0.6 and 0.7 (410 cases).

**Figure 5.**
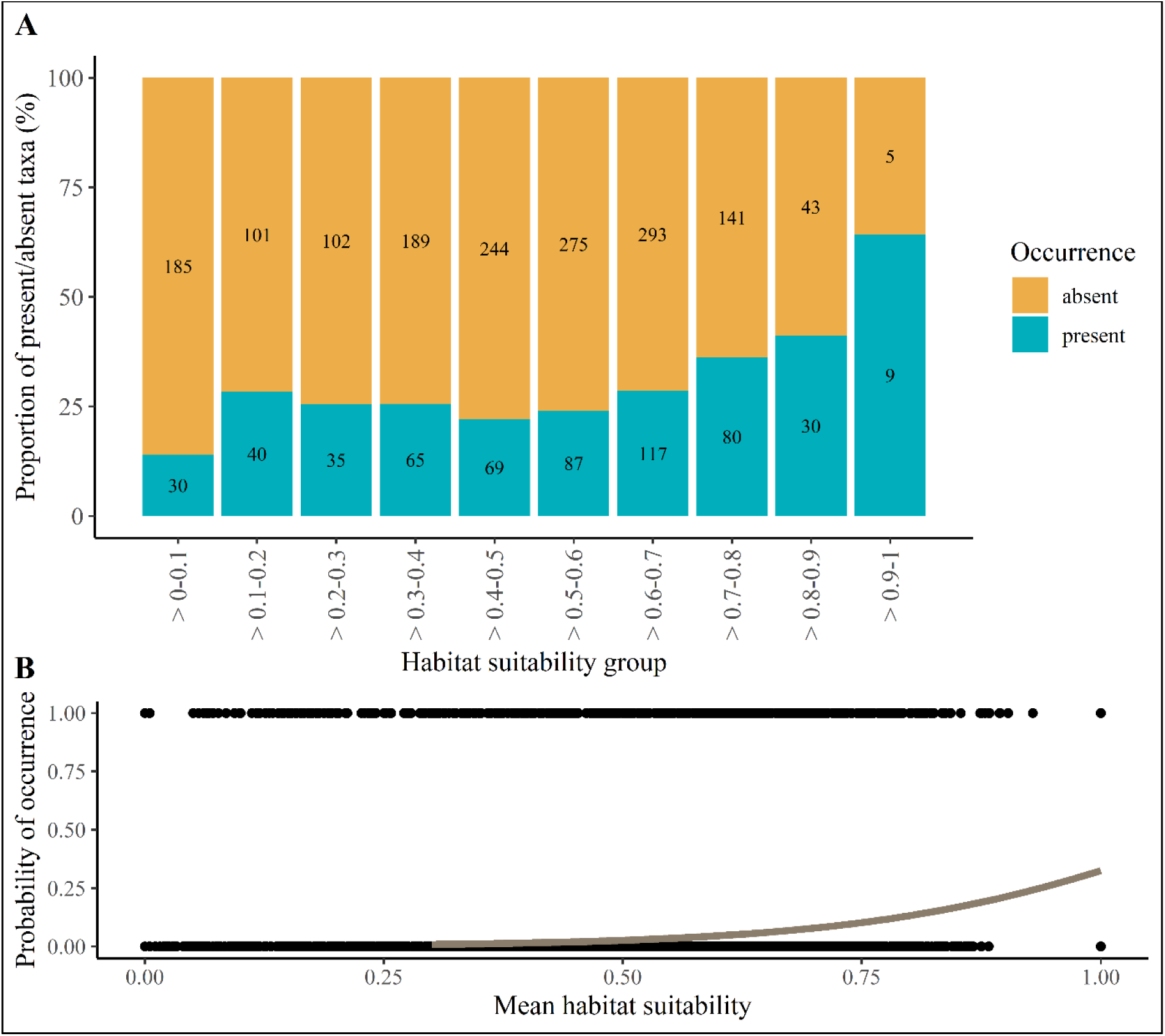
A: Proportion of present and absent taxa per habitat suitability group. The suitabilities were discretized into classes of 0.1, resulting in 10 groups. The number of ‘absent’ and ‘present’ cases is indicated in the corresponding bars. B: Logistic GLMM displaying the probability of occurrence in relation to mean habitat suitability (R² = 0.01).

Despite small differences in distributions, the mean habitat suitability is significantly higher for present taxa than absent taxa at unimpacted sites (Fig. 6, p < 0.05). A positive trend of the present taxa having higher suitabilities, as compared to absent taxa, is also observed at recently restored and mature sites. However, the difference at mature sites is only close to being statistically significant (p = 0.068), while at recently restored sites, it is not significant (p = 0.357).

**Figure 6.**
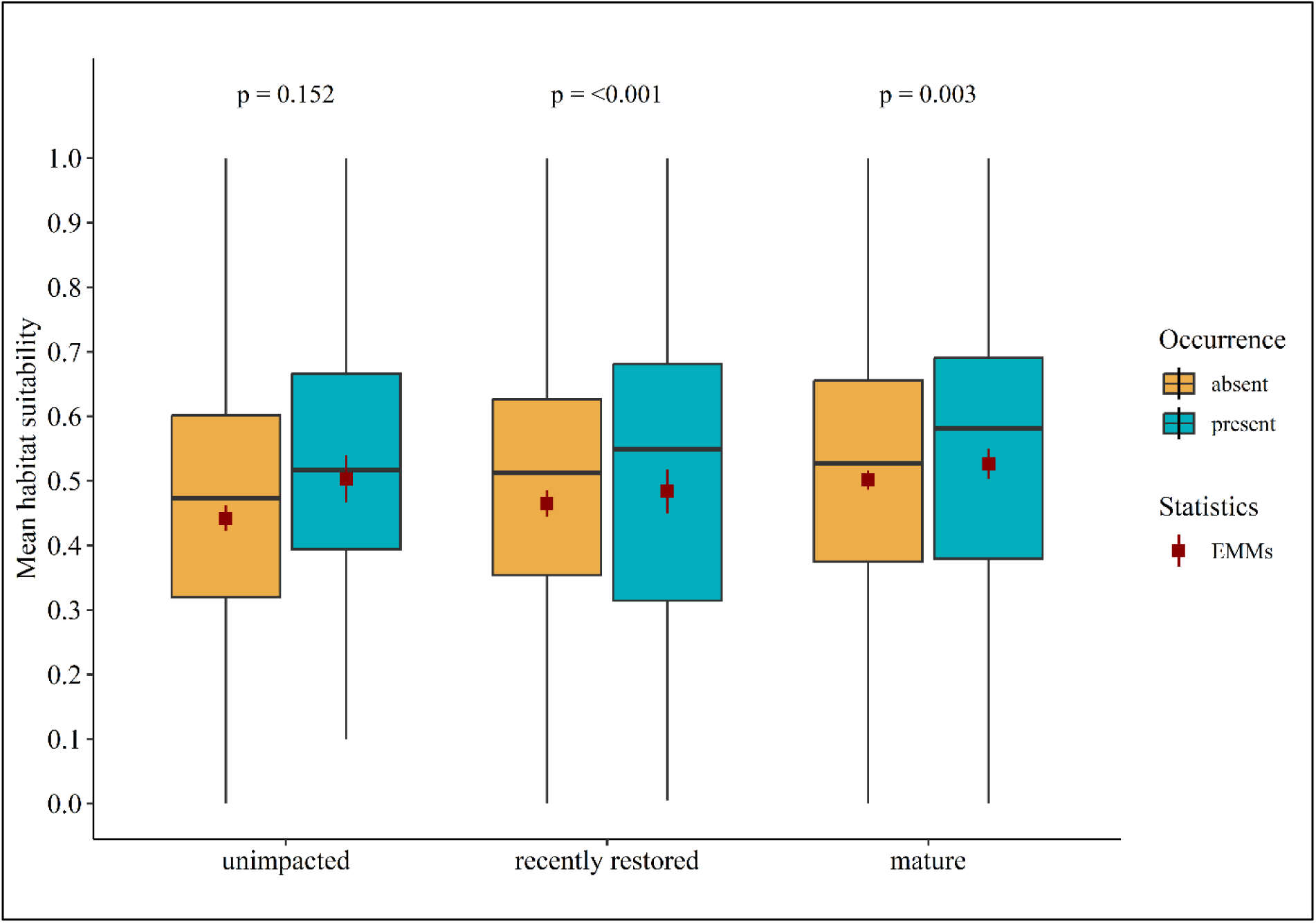
Boxplot of present and absent taxa and their mean habitat suitability. The difference between suitabilities of absent and present taxa were compared conducting a post-hoc test on estimated marginal means (EMMs). These were extracted from the corresponding generalised linear mixed model (GLMM), to account for different numbers of observations and to consider random effects. Estimated marginal means (EMMs) are represented by red boxes and corresponding error bars. Non-overlapping error bars indicate statistically significant differences (p < 0.05). For more detailed results on EMMs see Appendix S2, Figure S2.

### Biotic interaction filter

We determined the maximum Gower Similarity, i.e., trait overlap, between taxa that were and were not co-occurring per site. The proportion of present taxa is lower than that of absent taxa in all groups (Fig. 7A). The highest number of cases have a maximum trait overlap between 0.7 and 0.9. Modelled probability of occurrence predicts an overall increase along the gradient. However, the regression coefficient was not significantly different from zero (p= 0.068, Fig. 7B).

**Figure 7.**
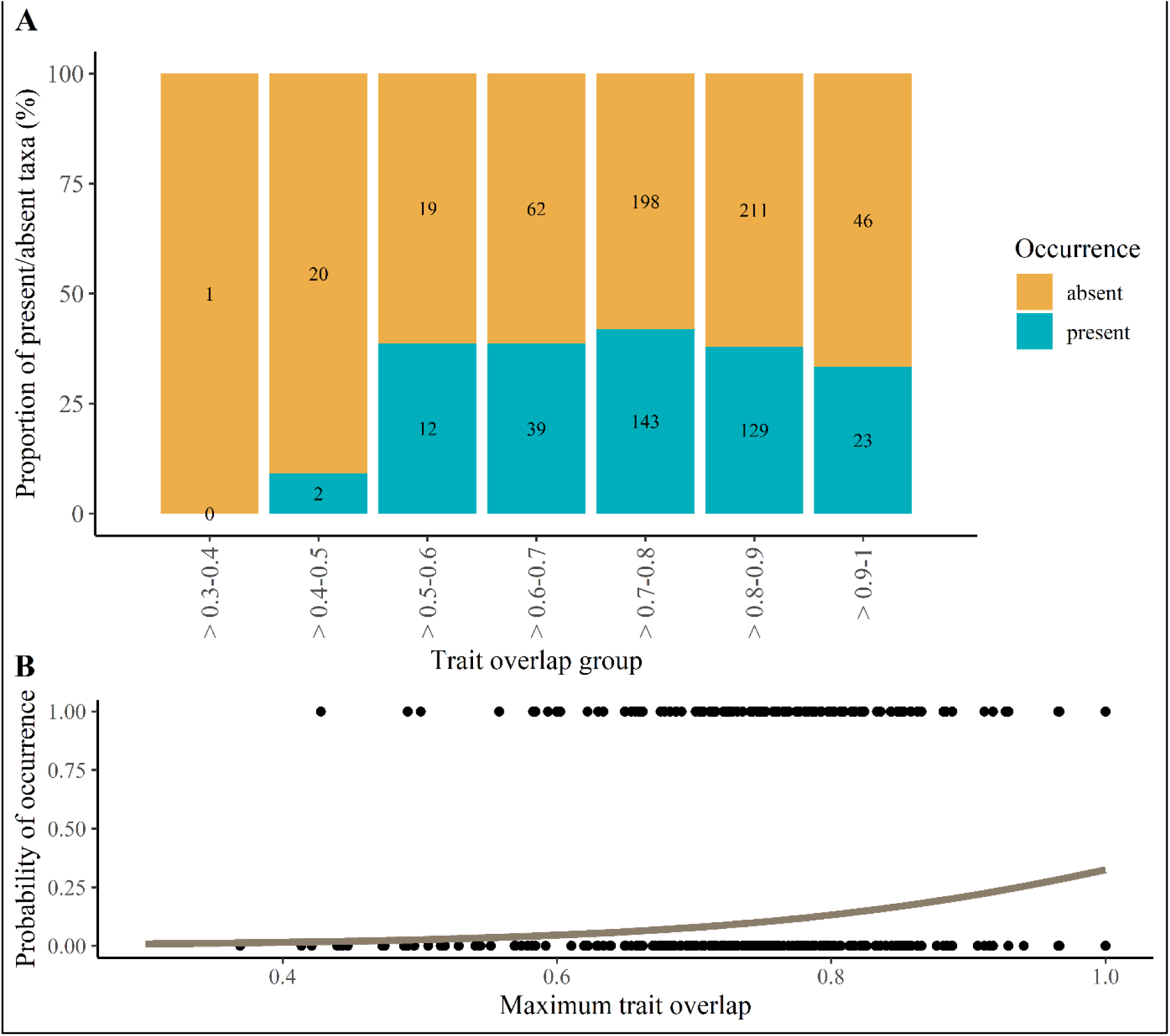
A: Proportion of present and absent taxa per trait overlap group. Trait overlaps were derived from the Gower similarities, calculated between co-occurring and not co-occurring taxa. Trait overlaps were discretized into classes of 0.1, resulting in 10 groups. The number of ‘absent’ and ‘present’ cases is indicated in the corresponding bars. B: Logistic GLMM displaying the probability of occurrence in relation to maximum trait suitability (R² = 0.01).

There are no significant differences in maximum trait overlap between co-occurring and not co-occurring taxa across the three groups of sites (Fig. 8). The median and EMM of the present taxa at unimpacted sites is slightly lower, however, this difference is not significant (p > 0.05). The spread of the data in the other groups is similar with no significant differences.

**Figure 8.**
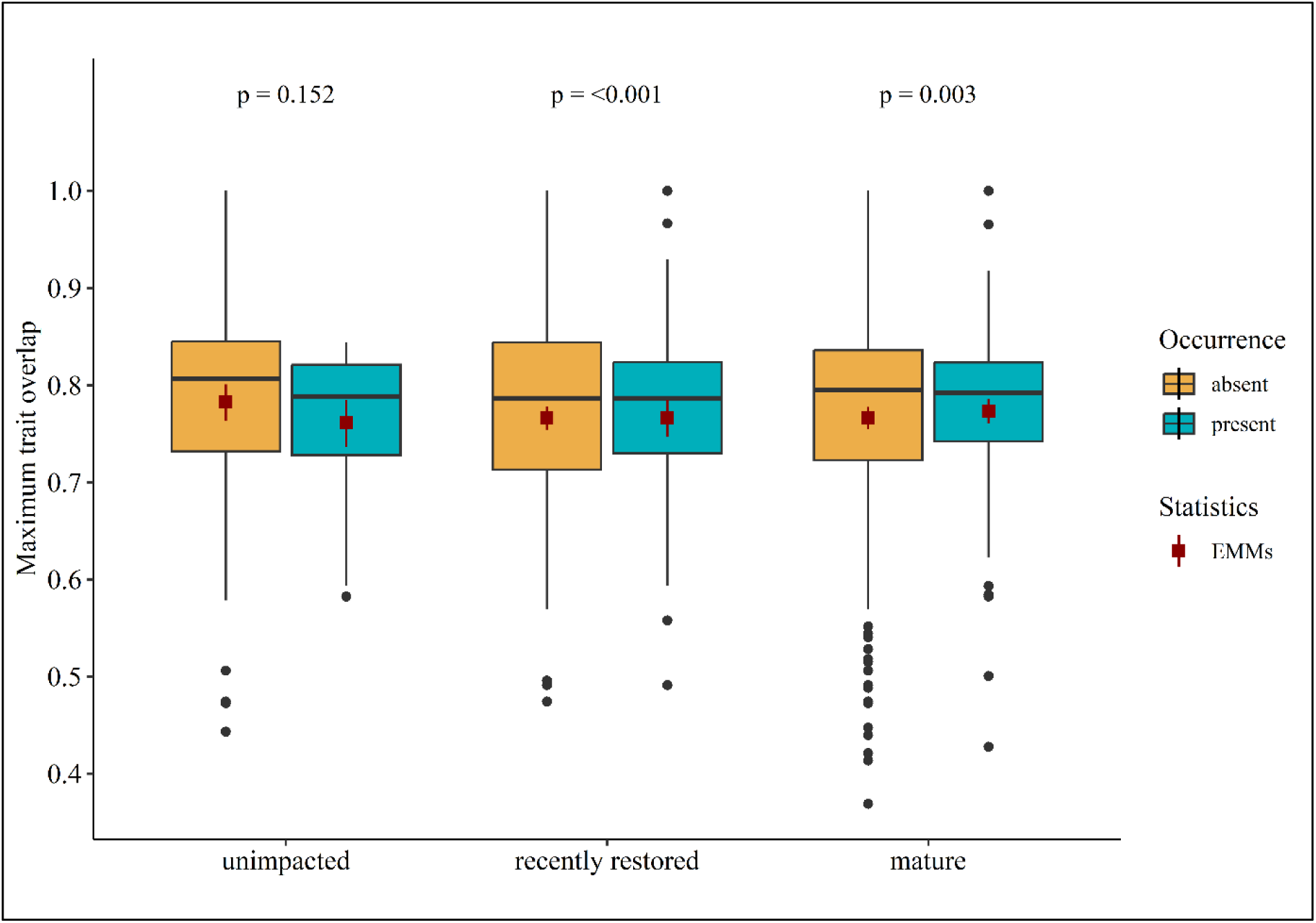
Boxplot of present and absent taxa and their maximum trait overlap. The difference between trait overlaps of absent and present taxa were compared conducting a post-hoc test on estimated marginal means (EMMs). These were extracted from the corresponding generalised linear mixed model (GLMM), to account for different numbers of observations and to consider random effects. Estimated marginal means (EMMs) are represented by red boxes and corresponding error bars. Non-overlapping error bars indicate statistically significant differences (p < 0.05). For more detailed results on EMMs see Appendix S2, Figure S3.

## Discussion

This study aimed to test the assumptions of the ARC with a recent and detailed dataset, comprising a total of 48 sites in the Boye catchment. More specifically, we challenged the assumption that the role of the colonisation filters, dispersal, environmental filtering and biotic interactions (competition) differs between stages of recovery. We first verified if the groups of sampling sites (‘unimpacted’, ‘recently restored’, ‘mature restored’) differ in their community composition, to justify further analysis. Indeed, the NMDS revealed that most communities of the recently restored sites (BOYohB224, LIEohBo, BOYuhHa, NATohBo) differ from those belonging to the other two groups. Additionally, the ANOSIM statistics suggested a moderate dissimilarity between groups which confirms our hypothesis (1) on differences between site groups. Similarly, Winking et al. (2014) found a significant difference between communities of recently restored sites (one to four years after restoration) and old restored sites (> 9 years after restoration) in the Boye catchment. The moderate outcome of the ANOSIM may be due to the generally low number of recently restored sites. Notably, the community at one of these sites (BOYohKi) was more similar to the mature Boye sites (BOY) than to the recently restored sites. However, this site is located in the middle of the catchment, in close proximity to the mouths of the Kirchschemmsbach and Haarbach and close downstream from old restored Boye sites, which may have resulted in a more rapid colonisation (de Donnová et al., 2022). Both, the sites LIEohBo and BOYohB224 share some characteristics with standing waters, which is a result from especially low flow velocities in large parts of the streams. Next to the recent restoration measures, this could be another explanation for their different community. The unimpacted and the mature sites did not differ from each other. However, sites from BRA (Brabecker Muehlenbach) were more similar to the Boye and QUAohBo, compared to the other two sites, belonging to the same group. The sites BRAob, BRAohBo and QUAohBo are located close together and have similar environmental conditions, thus offering similar habitats, and ensuring easy access for species from the same source populations. They are located directly upstream of the mature Boye sites, which was also reflected by their community similarity. This shows that the role of colonisation processes may not only differ between maturity stages but may also depend on the sites’ location within the stream network (Jones & Schmidt, 2018). Sites that are located close together and therefore well connected, have more similar communities, which are likely controlled by the same filters (Hughes, 2007).

Metacommunity assembly is controlled by different filters, acting in hierarchical order on species level, which were described by Lake et al. (2007). According to them, dispersal determines first, which species can reach the sites, environmental filters dictate whether those species can survive and biotic interactions, such as competition, allow only the strongest competitors to establish a population. To compare the role of the three filters between site groups, we used different metrics per factor, which we calculated for each combination of species and site. We then compared these per site group for present vs. absent species.

### Dispersal filter

We analysed the role of the dispersal filter, using the distances to source populations in the Boye catchment. To this end, we sampled 28 ‘source’ sites, in addition to the 20 ‘main’ sites, and determined per site and species the distance to the nearest upstream source population. The distance to source populations was deemed important for colonisation processes in previous studies (Sundermann et al., 2011; Tonkin et al., 2014). We therefore expected the probability of taxa occurrence to decrease with increasing distance to the nearest source (H2). Consequently, the proportion of taxa present per site should decrease. Our results support our hypothesis. We observed the highest proportion of present taxa to have their nearest source population within a distance of 1.8 km. Beyond this distance, the proportion of present taxa rapidly decreased. Notably, only a relatively small number of cases covered the distance between 0 and 0.5 km, while the highest number had a distance between 0.5 and 1.5 km to the nearest source. This was caused by our sampling design, in which we placed the potential ‘source’ sites 1 and 2 km upstream of each ‘main’ site. Upstream ‘source’ sites closer than 0.5 km were only present, if ‘main’ sites were located close together and thus serving as ‘source’ sites themselves for other ‘main’ sites. The matching of species’ occurrences between ‘main’ sites and upstream ‘source’ sites is most likely the result from mass effects of the upstream sites, i.e., the massive dispersal of individuals through drift or overland dispersal, which can be initiated by high competition pressure, flood events or mass hatching events (Leibold et al., 2004; Urban et al., 2006). In such cases, environmental conditions of the ‘main’ sites are less relevant for species to occur, due to continuous influx of new individuals from upstream. If larger distances need to be crossed, environmental conditions at the ‘main’ site gain importance for long-term establishment, as individuals leaving the site or not surviving cannot easily be replaced by immigrating specimens (Brederveld et al., 2011). A large proportion of absent taxa did not have an upstream source population available, which supports the finding that upstream population sources are a determining factor for species colonisation (Sundermann et al., 2011; Winking et al., 2014; Verdonschot & Verdonschot, 2023). However, a large number but rather low ratio of taxa, was occurring in individual sites despite the absence of a known and sampled source populations. These cases mainly include the unimpacted sites, for which no upstream samples were taken. Due to our sampling design, we may have missed additional source populations, less than 1km upstream. For our analysis, we have only considered upstream source populations. While we assumed this to be the easiest direction of dispersal for benthic invertebrates, dispersal can also be directed upstream, to compensate for aquatic passive downstream drift (Kopp et al., 2001). Especially, taxa with a flying adult stage (Ephemeroptera, Plecoptera, Trichoptera) can easily colonise from neighbouring streams and catchments. In fact, adult caddisflies were reported to fly distances of at least 1.5 km (Graham et al., 2017), although most individuals disperse close to their site of emergence (Collier & Smith, 1998). Passive aerial dispersal is also possible for hololimnic species, e.g., via phoresy (Figuerola & Green, 2002; Van Leeuwen et al., 2013). Hence, taxa could have entered the Boye system from a neighbouring stream network, e.g., the Rotbach or Schwarzbach (Enss et al., 2024). Both streams are in near-natural conditions and could act as potential source populations for the Boye system (Winking et al., 2016).

When comparing the different site groups, we expected that the dispersal filter would constrain taxa from occurring at the recently restored sites (H2). Thus, the present taxa should have a smaller distance to population sources than absent taxa. Distances to population sources differed significantly between present and absent taxa at the recently restored sites. While the same was true for the mature sites, the spread of distances for absent taxa was greater at recently restored sites. In addition, the estimated marginal means were further apart from each other at recently restored sites, compared to the mature sites. This indicates that a dispersal filter is acting on both, the recently restored and the mature sites, albeit stronger on the recently restored sites. Thus, the dispersal filter seem to have a larger role in the early recovery phase, which is in line with the predictions of Vos et al. (2023). In addition to the short time since restoration, this pattern might be increased by the fact that the recently restored sites are all located in the downstream parts of the catchment, further away from potential source sites, which might hamper fast colonization from upstream (compare Li et al., 2018). Taxa that were absent from recently restored sites had a much higher marginal mean than all others, but also distances of present taxa were slightly higher, compared to the mature sites. This indicates that source populations were not as easily available. The smaller difference between absent and present taxa indicates that the mature sites have already been colonised by most of the species occurring in the surroundings. In general, well-connected streams are more readily colonised, compared to unconnected ones (Hughes, 2007). At the unimpacted sites, the distance to the nearest source did not differ significantly between present and absent taxa. Additionally, the marginal mean of present taxa at these sites was higher than that of absent taxa. Upstream sources were unavailable for three of the unimpacted sites, hence the data only results from two sites that are close together, in the upstream part of the catchment. Although anthropogenically affected, these sites were never modified nor used as open sewer. Hence, their longitudinal position and near-natural conditions make them accessible and attractive for dispersing species.

### Environmental filter

Using the habitat suitability calculations, we tested the occurrence of taxa based on the presence of their preferred habitat. We expected that present taxa were characterised by a higher habitat suitability than absent taxa (H3). However, habitat suitabilities evenly ranged from 0.1 to 0.7. The proportion of present taxa increased only slightly towards the end of the gradient, similar to the probability of occurrence. However, at the same time, the number of cases per proportion decreased. Thus, while the general pattern fits our hypothesis, our test was not very strong. The comparison between groups of sampling sites showed that the habitat suitability of absent taxa was lower than that of present species at all sites. Nevertheless, the result was only significant for unimpacted sites. For the mature sites it was only close to significant. Therefore, we found indications that environmental filtering is mainly affecting the unimpacted and, to some degree also the mature sites. Depending on the extent of the restoration measures, existing habitats are removed or severely altered. At the start of the recovery phase, the available habitat is therefore limited. Such conditions allow mainly the occurrence of habitat generalists, resulting in medium habitat suitability values (Winking et al., 2014; Baumgartner & Robinson, 2017). As a result of riparian vegetation growth and organic matter transported from upstream, the habitat heterogeneity increases over time, which results in stronger environmental filtering (Barnes et al., 2013; Liu et al., 2021; Gillmann et al., 2023). Additionally, as previously mentioned, proximity to source populations leads to mass effects, which cause species to colonise less suitable habitats (Leibold et al., 2004). This is especially important in freshly restored streams, where habitats are less readily available, since we observed no significant difference between habitat suitabilities of absent and present taxa. On the other hand, at the mature and unimpacted sites, habitats may have stabilized, facilitating a stable and more specialized community, which translates into higher habitat suitability, especially at unimpacted sites (Gillmann et al., 2023). However, if sites are close together, mature sites may, too, be affected by mass effects from upstream.

In general, our data showed a large mismatch between expected and observed taxa, based on the habitat suitability. This problem was also identified previously in habitat suitability modelling (Brantschen et al., 2024). Presence/absence data might be less indicative for environmental filtering than abundance data because species may be found at less suitable habitats but in lower abundance (Johnson & Vaughn, 1995). Highly suitable habitats would in turn be populated in great numbers. This could also explain the small differences between present and absent taxa per habitat suitability group. Many taxa are not restricted to only one type of substrate, although they may favour some over others. The database freshwaterecology.info accounts for this by defining different affinities to every possible type of substrate. Hence, if a taxon is assigned a low affinity for a certain substrate type, present at the site, the habitat suitability will be lower than if a high affinity for the same substrate is assigned. The taxas’ potential to also occur in habitats, not reported by freshwaterecology.info could have also led to the mismatch between the assumed suitability and actual occurrence. In addition, sites within a small catchment, such as addressed here, may not be large enough to exhibit distinct patterns of present and absent species.

### Competition filter

We assessed the role of competition for community assembly, using the Gower similarity index as a measure for trait overlap. To determine if the biotic interaction filter influences the communities of the different site groups, we compared the maximum trait overlap between co-occurring and not co-occurring taxa. As a consequence of competition, we expected the trait overlap between co-occurring taxa to be low, while that of taxa not co-occurring should be high. Accordingly, the proportion of present taxa should decrease with increasing trait overlap.

In contrast to our expectation, the proportion of present taxa was never higher than that of absent taxa, for none of the trait overlap groups. Instead, the proportion of present taxa was always around 40 % across all trait overlap groups. The comparison between site groups did not reveal any significant differences. In contrast to our expectation, the probability of occurrence increased with increasing niche overlap. Only at the unimpacted sites, we observed the trend that present taxa had a lower trait overlap, compared to absent taxa. These results leave do not conclusively resolve our hypothesis since variations in trait overlap were not large enough to show a clear pattern.

Quantifying competition between species is difficult in field studies since trait overlap does not rule out co-existence. The higher similarity between traits of taxa within the same community may result from habitat stabilisation. This aligns with more stable communities that likely prefer similar environmental conditions (Gillmann et al., 2023). Previous studies have shown that the mean trait overlap within restored communities increases with time since restoration, which was defined as an increase in potential interspecific competition (Gillmann et al., 2024). While species with similar traits are more likely to compete with each other, Schlenker et al. (2024) demonstrated that trophic similarity increases over time, indicating that the existing species occupy similar trophic niches. Regional co-existence relies on the local availability of suitable habitats for the competitors. If co-occurring species with high trait overlap are equally good competitors, only the comparison of their abundance will inform, whether one of the populations is suppressed. This could be particularly important on the microhabitat scale. Different patches within the same sampling site can be occupied by different species with similar preferences (Moser & Minshall, 1996). As described by Johnson & Vaughn (1995), densities can identify successful competitors within these patches. Fast reproduction may then be of greater importance than the preferences itself, in order to quickly take up the available niche space. Overall, regional competitive exclusion was previously shown to be a slow process, which can lead to changes in species distributions even if the environmental conditions remain stable (Yackulic, 2017). Thus, trends of increasing and decreasing abundances of possible competitors over time, should be considered for the assessment of competitive interactions within communities.

The lack in differences between trait overlap of co-occurring and not co-occurring taxa could also result from not taking the dispersal filter into account in the first place. From our results, regarding the distance to source populations, we have observed that in many cases, an upstream source population is missing. Other source populations are far away (> 2 km). Hence, only taking taxa with upstream source population into account, could lead to a better explanation of species absences. Species that are not able to reach a respective site, cannot compete with occurring species, making their degree of niche overlap negligible.

## Conclusions

The process of recolonisation is key for the recovery of restored streams. In line with our first hypothesis, the recently restored sites differed from all other sites (H1). Hence, the recolonisation process takes time and is limited by the dispersal filter during early phases of recovery. Early communities consist of species that occur in the close surroundings of the restored sites and therefore strongly depend on nearby source populations (H2). Environmental filtering is especially important in well-developed habitats, where specialised species have established a stable community (H3). We did not observe competition to be important for community assembly (H4). However, competitive exclusion could be important on the microhabitat scale and become more apparent when abundances are taken into account.

Our findings support the assumption that the role of the dispersal and environmental filter differs between different phases of recovery, as proposed by the ARC. Thus, a good connectivity to source populations and support to develop complex habitats during succession are a promising foundation for restored streams to support a diverse benthic invertebrate community. Our results could not support the expected increase in importance of the biotic filter. Thus, future research should be directed at more detailed investigations of interspecific interactions.

## Acknowledgements

We thank the student assistants, Jannis Budke, Ronja Finke, Carolin Klancicar, Mona Roschewski and Nele Wittmeier, for their help with field sampling and sample processing. A special thanks to the Aquatic Ecosystems research group at the University of Duisburg-Essen, for performing the DNA metabarcoding of our samples. We appreciate the support of Wim Kaijser regarding statistical analyses. Finally, we are grateful for the collaboration with the Department for Systems Analysis, Integrated Assessment and Modelling at the Swiss Federal Institute of Aquatic Science and Technology (Eawag). This work results from the CRC RESIST, which was funded by the German Research Foundation (Deutsche Forschungsgemeinschaft, DFG) – CRC 1439 – project number: 426547801.

