## Appendix S1 for "Contributions of source populations, habitat suitability and trait overlap to benthic invertebrate community assembly in restored urban streams"

Appendix S1: Additional information for data processing

Table S1. Eight microhabitat preferences of benthic invertebrates and the corresponding habitat types, as defined in freshwaterecology.info (Schmidt-Kloiber & Hering, 2015)

| Substrate preference | Explanation |
| --- | --- |
| Argyllal | Silt, loam, clay (grain size < 0.063 mm) |
| Pelal | Mud (grain size < 0.063 mm) |
| Psammal | Sand (grain size 0.063 – 2 mm) |
| Akal | Fine to medium-sized gravel (grain size 0.2 – 2 cm) |
| Lithal | Coarse gravel, stones, cobbles, boulders, bedrock  (grain size > 2 cm) |
| Phytal | Algae, mosses, macrophytes |
| POM | Coarse (CPOM) and fine particulate organic matter (FPOM) |
| other | Other substrates |

Table S2. LAWA water quality standards for chemical parameters, and the SI (DEV 1992, 2004; LAWA 1998); published in Bernatowicz et al., 2009.

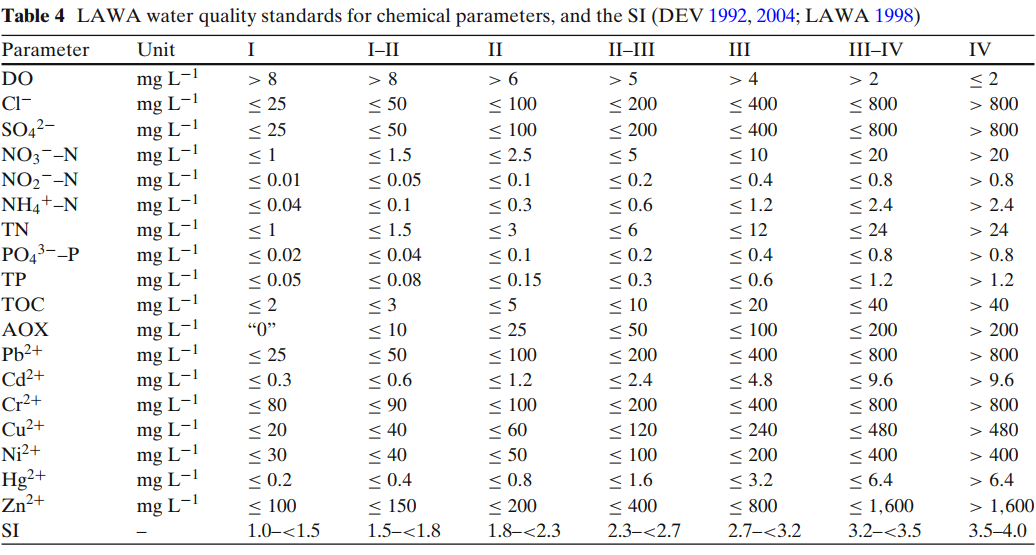

Table S3 Borders defined to suit the categories of the species’ preferences.

| Trait | Weight | Borders | Source |
| --- | --- | --- | --- |
| KLIWA index | 0, 1, 2, 3, 4, 6, 10 | 1, 3, 5, 7, ,9 11, 100 | Sundermann et al. (2022) |
| Saprobic value | 4, 8, 16 | 1, 3, 4 | German saprobic index |
| Flow velocity | - | 0, 0.01, 0.02, 0.05, 0.06, 0.09, 0.1, 0.15, 0.16, 0.25 | STOWA, Verberk et al. (2012) |
