## Appendix S2 for "Contributions of source populations, habitat suitability and trait overlap to benthic invertebrate community assembly in restored urban streams"

Appendix S2. Results of Estimated Marginal Means

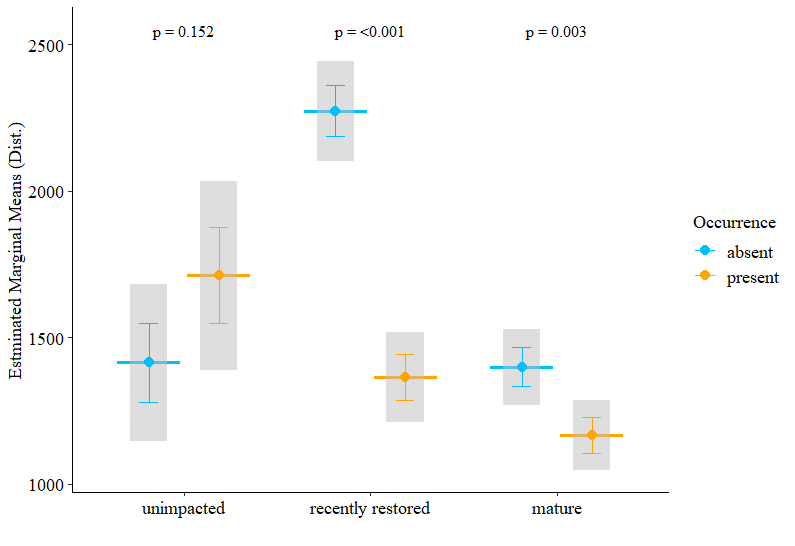

Figure S1. Estimated Marginal Means (EMMs) extracted from GLMM with ‘distance to source’ as response, ‘occurrence’ and ‘site group’ as fixed and ‘species’ as random effect. Grey vertical bars display the 95% confidence intervals. Horizontal bars depict the EMMs. The difference is statistically different, if standard error bars do not overlap (p< 0.05).

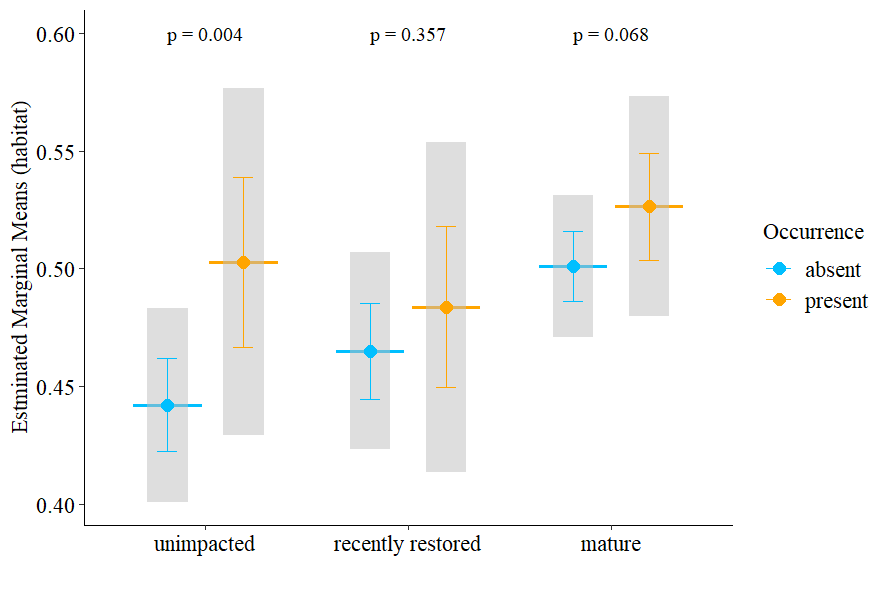

Figure S2. Estimated Marginal Means (EMMs) extracted from GLMM with ‘mean habitat suitability’ as response, ‘occurrence’ and ‘site group’ as fixed and ‘species’ as random effect. Grey vertical bars display the 95% confidence intervals. Horizontal bars depict the EMMs. The difference is statistically different, if standard error bars do not overlap (p< 0.05).

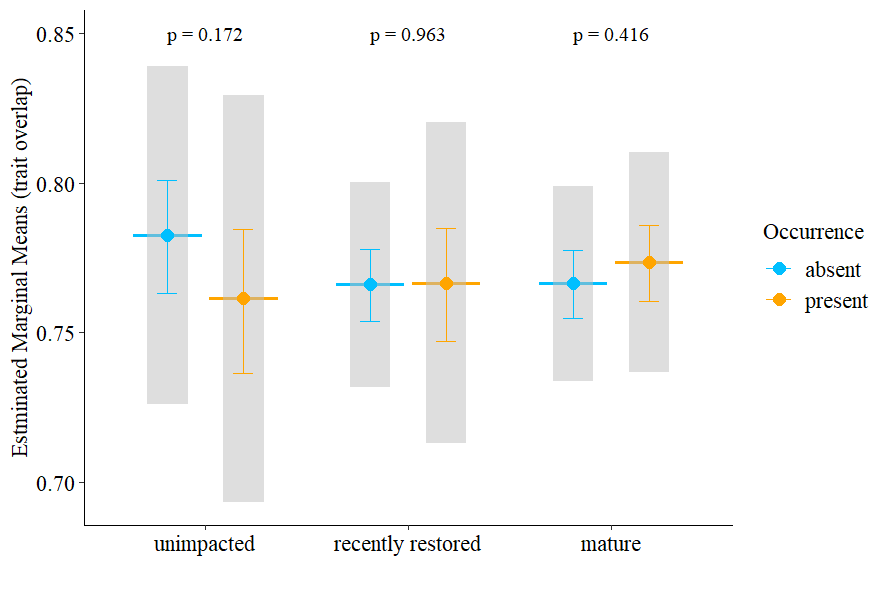

Figure S3. Estimated Marginal Means (EMMs) extracted from GLMM with ‘trait overlap’ as response, ‘occurrence’ and ‘site group’ as fixed and ‘species’ as random effect. Grey vertical bars display the 95% confidence intervals. Horizontal bars depict the EMMs. The difference is statistically different, if standard error bars do not overlap (p< 0.05).
